# Solvation Dynamics of a Single Triglyceride as a Function of its Chain Length

**DOI:** 10.1101/2022.03.28.486057

**Authors:** Sukriti Sacher, Arjun Ray

## Abstract

Triglycerides (TG) are transported packaged inside lypophillic particles. Several lipid exchange/transfer proteins interact with these lipoproteins and facilitate lipid exchange amongst lipoproteins, to maintain a constant flux in RCT. During this process, these neutral lipids are inadvertently exposed to the bulk water. Previous studies have elucidated the behavior of triglycerides in the bulk (on the surface of bilayer or inside a lipid droplet). However, isolated TGs during lipid exchange behave differently than when in bulk, due to an increased exposure to water. We studied the solvation dynamics of a single TG in a polar (water) and a non-polar (cyclohexane) medium to elucidate it’s solvated structure while drawing parallels with its structural organization in bulk (lipid droplet). We also examine the role of acyl chain length and it’s contribution to the free energy of solvation. Finally, we have established the predominant conformation of TG in water and cyclohexane and discuss the thermodynamics for such a preference.

## Introduction

Triglycerides (TGs) represent an important source of cellular energy. They are the predominant form in which fatty acids are stored and transported ^1^. Beyond their role in energy homeostasis, they form an intrinsic component of cell membranes, participating in metabolic processes that involve cell stimulation, transformation and metastasis ^2^. A condensation product between three fatty acids and a glycerol, TGs encompass a heterogeneous group that vary in the length of their hydrocarbon chains and unsaturation. Depending on their fatty acid composition, they can be made up of long chain fatty acids (LCFA) (14 or more carbons), predominant in fish and fish oil ^3^, medium chain fatty acids (MCFA) (6-12 carbons) that is prevalent in coconut and palm oils ^4^ and short chain fatty acids (SCFA) (*<* 6 carbons) which are mainly produced by gut microflora in addition to being an important constituent of cow milk ^4;5^. Owing to their highly reduced state, hydrophobicity and limited water solubility ^6^, they are stored as lipid droplets in cells and are transported within specialized lipoprotein vehicles while in plasma. TG-rich lipoproteins (TRLs) are mainly secreted by intestinal enterocytes and hepatocytes. These mainly constitute chylomicrons, Very Low-Density Lipoprotein (VLDL), its remnants and Low-Density Lipoprotein (LDL). The plasma concentration, structure and dynamic composition of these lipoproteins is of immense biological importance as their dysregulation is a major causal factor for atherosclerotic cardiovascular disease (ASCVD) ^7;8^. Hypertriglyceridemia, caused due to an increase in TRLs in plasma has been shown to contribute to intimal cholesterol deposition, activation and enhancement of pro-inflammatory and pro-coagulant pathways as well as concomitant changes in high density lipoprotein (HDL) ^9^. These changes are associated with a substantially increased long term total mortality and risk for cardiovascular events ^8;10^.

Understanding the fate of TRLs and their metabolic effect, requires an extensive understanding of its components. Lipoprotein shape, surface structure, and metabolic fate has been shown to be determined by the physical state of its core which in turn is a reflection of its composition ^11^. Several in-silico studies have attempted to model and characterize the TG-rich core in these lipoproteins. Hall et al. provided a comprehensive understanding of TG packing as well as phase transitions in TG aggregates, highlighting the dynamic properties of mixing, interdigitation and diffusion in a pure TG melt ^12^. Studies focused on the surface of LD-mimicking systems have emphasized that TGs intercalate into phospholipid (PL) tails while retaining their core-like disorder. This drastically influences the molecular properties of the surface of the lipid monolayer that surrounds LDs ^13^. Several other studies have observed the tendency of TGs to partition onto the surface of the droplet in addition to the hydrophobic core forming a disordered, isotropic, aggregate under the PL layer ^14^. The presence of TG on the surface has also been shown to alter the mechanical property of monolayer drastically by decreasing its bending elasticity which may have important implications in cellular transformation and metastasis ^15^. Recently, it has also been shown that the surface TG is ordered like the surface phospholipids with its glycerol group oriented towards water and acyl tails pointing towards the center of the bilayer ^16^. The ability of glycerol group to participate in H-bonds stabilizes this orientation and also leads to hydration of LD core ^16^.

While several studies have characterized the bulk properties of TGs in LD core or surface, no studies to date have provided an atomic description of the solvation of an isolated TG, while simultaneously drawing relations to the structural organization it orchestrates in the bulk. Solvation of a single molecule, otherwise considered to be a complex phenomenon, is influenced by the atomic structure of the molecule that facilitates non-bonded interactions with the solvent. An interplay between these forces together, governs solute properties such as solubility, diffusion and aggregation. These properties in turn influence the other bulk properties ^17;18;19^. Solvation behavior of an isolated TG is also of immense importance in mechanisms of exchange and transfer by proteins. These lipid transfer proteins such as microsomal triglyceride transfer protein (MTP), cholesterol ester transfer protein (CETP), selectively interact and access these neutral lipids (one at a time) from the core of lipoproteins and exchange it with other lipoproteins ^20;21^. The hydrophobic interactions between residues of the protein that interact with TG, shield it from bulk water during it’s transit ^22;20^. These protective hydrophobic sites in these neutral lipid LTPs, mimic TG’s native non-polar environment, while it is exchanged between vesicles in aqueous medium. An in-depth understanding of this transfer process that encompasses thermodynamics and kinetics requires an understanding of the behavior of TG when immersed in water.

In this work, we have established an atomic scale description of the solvation structure of an isolated triglyceride using molecular dynamics simulation. We have characterized the thermodynamics of the process of solvation as a function of the acyl chain length by considering three different lengths designated as long (18:0/18:0/18:0), medium (10:0/10:0/10:0) and short (4:0/4:0/4:0) (Supplementary Figure 1 & 2) in a polar (water) and a non-polar (cyclohexane) solvent. We have also established the stable TG conformation in each solvent type and have attempted to characterize the inter-conversion between these conformers that the solute-solvent interactions facilitate. We highlight the differences between the structural organization of a single TG when surrounded by other TGs as in a droplet as opposed to when surrounded by water. The observed results provide an insight into structure and dynamics of TG at the lipid-water interface during its partitioning to the surface of bilayer or during its transit through the lipid exchange proteins.

## Results and Discussion

On being immersed in a solvent, the organization of solvent molecules around TG were first evaluated using the radial distribution functions [RDFs, *g(r)* between functional groups of a TG (solute) and solvent molecules in terms of their site-site pair correlations. A solvent *g(r)* profile is a function of the density of the solvent molecules at a distance r from a specified point/atom/group normalized by the bulk density of the solvent. Figure 1A illustrates the *g(r)* profile between the center of mass (COM) of TG and water molecules. We observed a broad, less intense peak in case of long TG at ∼0.45 nm which gets sharper as the acyl chain length decreased. This indicated that long TG keeps water molecules further apart than medium and short TG while forming its first solvation layer. Integration of the COM_TG_ – water *g(r)*) profile upto the first peak minimum yielded ∼29 water molecules for long ∼38 and ∼45 molecules for medium and short TG respectively. In case of cyclohexane, it was observed that in each case, irrespective of the acyl chain length, three peaks were obtained representing three solvation layers, however, the intensity of each of these peaks declined subsequently (Figure 1B). Presence of multiple solvation layers is indicative of hydrophobic forces higher in magnitude that enable TG to keep cyclohexane molecules closer.

**Figure 1:**
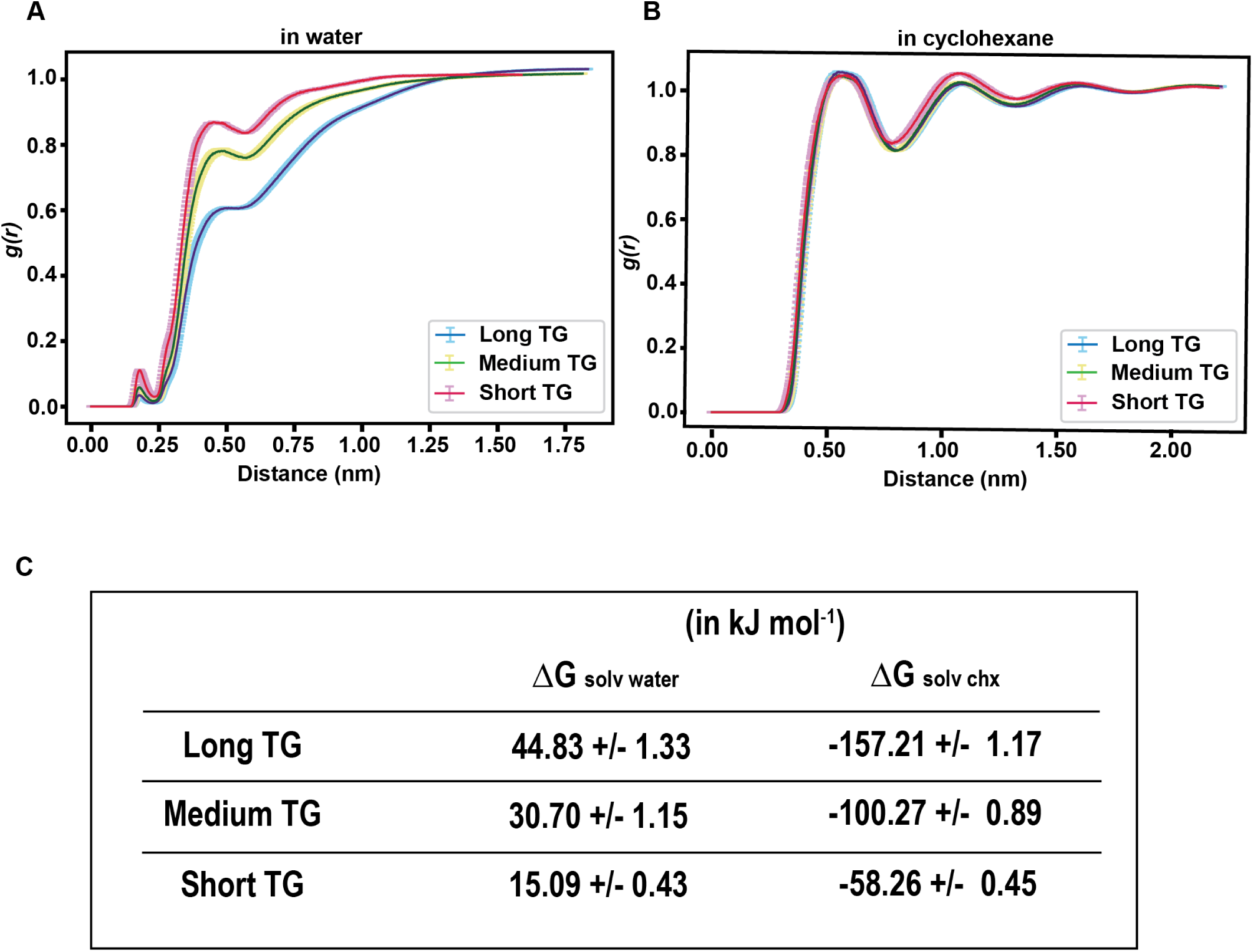
Solvation dynamics of a single triglyceride. **A**. Radial distribution profile of water molecules around the center of mass of the three TGs varying in acyl chain length designated as long (indigo), medium (darkgreen) and short TG (crimson). Solid lines represent the mean values while error bars depict the absolute deviation from the mean calculated across three replicates of simulation. **B**. Radial distribution profile of water molecules around the center of mass of the three TGs. **C**.Solvation free energies of the three TGs calculated by thermodynamic integration in water as well as cyclohexane.

We also observed a sharp but less intense peak at ∼0.20 nm in the *g(r)* profile of all the three TGs in water (Figure 1A), but not in cyclohexane (Figure 1B). We assumed this to be a result of H-bond interaction prevalent in water but not in cyclohexane. In order to test our hypothesis, we selectively computed the *g(r)* profile of H-atoms of water around the carbonyl oxygen of acyl tails of TG. A peak at ∼0.20 nm was observed and it was found to overlap in all the three cases of acyl chain length (Supplementary Figure 3), therefore we concluded that the H-bond attraction between carbonyl oxygen of acyl tails pull a few water molecules closer to TG. It may be noted that the oxygen atom from the glycerol group also participates in a few H-bond interactions with water (Supplementary Figure 4). But the intensity of this interaction is less than that of the carbonyl oxygen. In order to elucidate the contribution of acyl tail and glycerol group in TG solvation, we computed the radial distribution profiles of solvent molecules around these functional groups separately as well (Supplementary Figure 4). In cyclohexane, the profiles for acyl tails and glycerol appeared similar and were found to closely overlap, implying that the distribution of cyclohexane molecules around TG is unaffected by functional group or chain length. However, in water, a less intense peak was observed in case of both acyl tail and glycerol at ∼0.20 nm. This peak was sharper in case of acyl chain than glycerol and is indicative of the hydrophilic nature imparted by the O atom in both these groups. This was followed by a broad peak that dampened as the acyl chain length increased. The difference in intensity of the peak can be ascribed to the increased hydrophobicity of the acyl chain as its length increases. Therefore, we concluded that the number of water molecules in the first solvation layer as well as the distance at which this hydration sphere is found is a function of the hydrophobic character of TG, and this is influenced by the number of carbon molecules that constitute its acyl chains.

Our findings of the structural properties of a solvated TG were further supported by the computation of free energy of solvation using the thermodynamic integration method (TI) ^23;24^ (Figure1C). Free energy of solvation is defined as the amount of energy required to introduce the solute into the solvent, and can be used to evaluate the relative stability of interactions between these two components ^25^. The TI method utilizes the *λ*-dependence of the Hamiltonian *H*, which in turn is a function of the coupling parameter *λ* and defines a pathway that connects the solvated state with the unsolvated state (See Methods for more details). We evaluated the ensemble average at a number of discrete *λ*-points by performing separate simulations for each chosen *λ*-point. The sum of free energy changes of each of these discrete points, describes the total free energy change from *λ* = 0 (when the solute-solvent interactions are uncoupled) to *λ* = 1 (when solute interacts with the solvent). Considering the free energy of solvation computes the energy used to decouple the solute from the solvent, therefore, the results obtained are actually a reverse of the solvation energy. Our computation indicated that the ΔG_solv_water for all the three TGs was positive, indicative of the unfavorable environment provided by water. However, negative values were obtained in case of ΔG_solv_cyclohexane (Figure 1D). Moreover, the amplitude of the solvation free energy was observed to be a function of acyl chain length.

We furthered this idea by calculating the free energy of solvation of symmetric TGs with acyl chain lengths ranging between four carbons to eighteen carbons, encompassing an array of short, medium and long chain TGs. Supplementary Figure 5A & B depicts the relative free energy differences for each interval of *λ* (between neighboring Hamiltonians) for the eight TGs in water and cyclohexane, respectively. Moreover, the magnitude of ΔG_solv_water & ΔG_solv_cyclohexane is directly proportional to the number of carbons in acyl chain (Supplementary Figure 5C-E). It is evident that cyclohexane provides a favorable environment for van der Waal interactions between the acyl tails and itself and this interaction is more intense in TGs with longer hydrocarbon chains. The opposite is true for water, where the longer acyl tail exerts an opposite force inhibiting its dissolution.

Next, we wanted to evaluate the effect of hydration on a single TG. In a lipid droplet core (bulk), TG was previously shown to exist in six conformations assigned on the basis of the orientation of the acyl tails about the central carbon (Figure 2A) ^13^. Utilizing this same description, we discretized our simulation frames into these six states, namely stacker, trident, T, hand, fork and chair. Those orientations that deviated largely from any of these six conformations were assigned as ‘Other’ (See Methods & Supplementary figure 6-9 for details). The frequency of each conformation throughout the simulation was calculated to assess the propensity of TG to exist in a particular form in either water or cyclohexane as opposed to a lipid droplet suspended in water. Figure 2B shows that long TG preferred to exist as trident in water, T in cyclohexane and stacker in the lipid core. While, medium and short TG, either water solvated or present inside a droplet, preferred to exist as T (Figure 2C & D).

**Figure 2:**
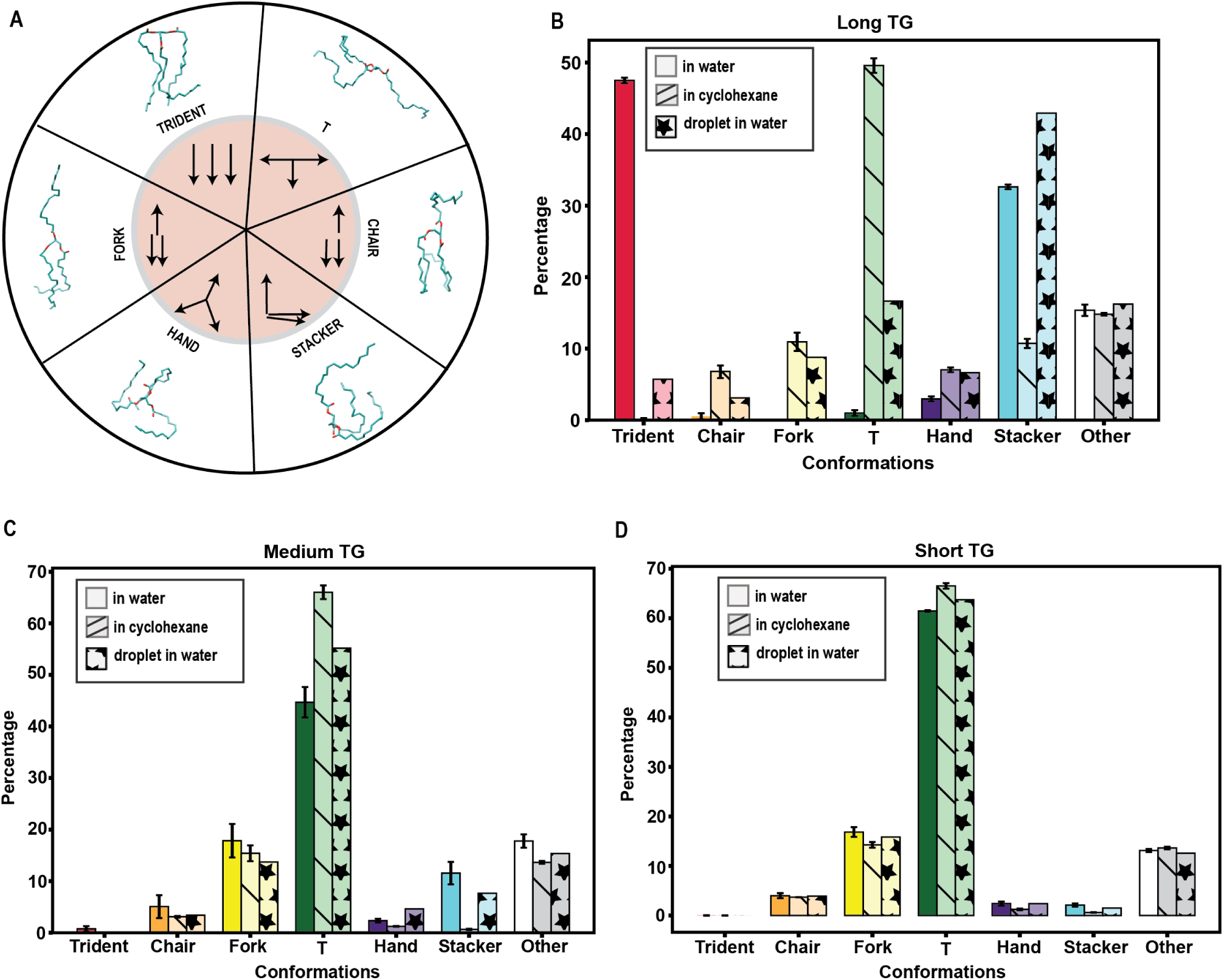
Distribution of the different conformations of a Triglyceride. **A**. The six conformations that a triglyceride molecule can exist in. Panel shows a schematic representation along with a representative image for each of the six conformations. (Adapted from Backle et al. ^13^. Frequency of each of the six conformation of a TG, either **B**. long **C**. medium or **D**. short when solvated, or packaged in a lipid droplet core.

A similar preference for stacker conformation when TGs are packaged in a droplet and for T when surrounded by a non-polar environment has been previously highlighted in studies that studied a lipid droplet in water and a lipid droplet sandwiched between a phospholipid bilayer, respectively ^13^. Therefore, the prevalence of single TG to exist in a particular conformation in these two solvents, is in an agreement with its previously observed property in bulk. However, we observed that a single TG when hydrated largely exists in the trident form. While, there are minor differences between the way acyl tails are oriented in each of the six conformations, prevalence of a particular conformation warrants its stability in the medium.

Considering we observed that the solvation of TG is entirely dependent on the number of hydrocarbons, we presumed that the stability of a particular conformation in water would be dependent on the structural organization of water around the acyl tails. Therefore, we computed the number of water molecules (within a shell radius of 0.5nm) that surround the three acyl tails when a TG takes on a particular conformation. A shell radius of 0.5 nm was decided based on the rdf (Figure 1A) which peaks around 0.45nm. Although, we did not perceive any major differences in the affinity of water molecules to distribute around the six conformations, we did observe that the trident and stacker form of the long TG, were surrounded by fewer water molecules on an average compared to the other conformations (Supplementary Figure 10). This was also confirmed from the *g(r)* of COM of TG existing in each of these six conformations (Supplementary Figure 11) It may be noted that the COM of acyl tails may overlap entirely for certain conformations such as chair and fork and may lie very close in case of others. Therefore, the reordering of water molecules around the different conformations may not be distinguishable by the *g(r)*.

Despite the propensity of long TG to exist as stacker and that of medium and short TG to exist as T in water, we observed that these conformations were found to fluctuate between the other five conformations throughout the simulations (Supplementary Figure 12). Therefore, we evaluated if there was a directionality to this inter-conversion. For every conformation in our simulation, the frequency of it being succeeded by every other conformation was counted and has been tabulated in Supplementary Table 1, based on which a weighted network was constructed (Figure 3). This network represents the inter-conversion of conformations and the propensity of this inter-conversion is depicted by the thickness of each edge that joins two nodes. Figure 3A clearly depicts that stacker makes the maximum connections. Therefore, for long TG in water, each conformation has a propensity to convert to stacker, such that all paths no matter the starting or ending conformation, go through stacker. In other words, stacker forms the intermediate state for most of the inter-conversions observed. It may also be noted that propensity of conversion between fork ← stacker, fork ← T & inter-conversion of chair ↔ T is very similar. Additionally, stacker also converts to trident which is the form in which long TG prefers to exist (Figure 2A). It may be emphasized that this network only depicts the propensity of inter-conversion and the size of the node is not indicative of the prevalence of the conformation. In other words, trident form may still be the most predominant form for long TG, which is determined by the total number of frames in which TG exists as a trident, but stacker is the favoured intermediate. This means, every time there is a switch in conformation from trident or another conformation, to any of the other four conformations, it goes through stacker.

**Figure 3:**
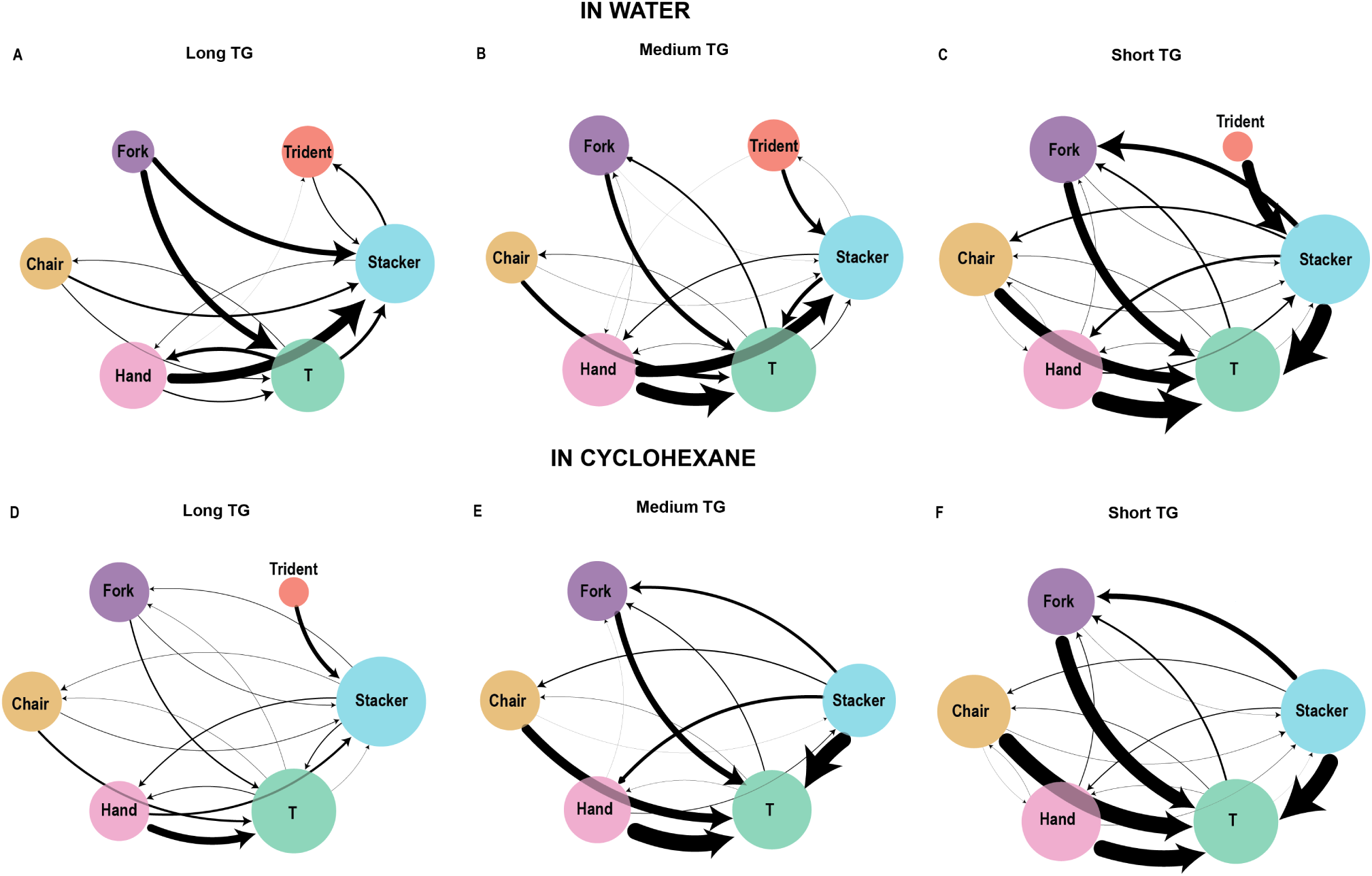
Inter-conversion of triglyceride conformations. A weighted network showing the inter-conversion between the six conformations. Panel A & D depict the inter-conversion of long TG, while B & E depict the inter-conversion of medium TG in water and cyclohexane respectively. Inter-conversion of short TG in water and cyclohexane is depicted in panel E & F respectively. Each node in the network represents one of the six conformations as indicated in the figure and its size is proportional to the number of connections with other nodes. Nodes are connected to other nodes through edges which are directional such that the direction of arrow points towards the conformation after conversion. The thickness of the edges are proportional to the frequency of conversion observed between the two conformations that they connect.

Medium and short TG in water as well as long TG in cyclohexane show a similar trend (Figure 3B-D). In all three of these networks, stacker and T are both the most popular nodes making a similar number of connections, but the propensity of conversion to T is much higher than stacker. T was observed to convert to fork, chair and even stacker, albeit, less favorably. Another interesting trend is that stacker is the preferred intermediate for conversion from trident to any other form which is evident from a single thick edge emanating from trident to stacker and a complete absence of connections to the other nodes. Figure 3E & F depict the network of inter-conversion amongst conformations of medium and short TG in cyclohexane. In this case too, T has the highest number of connections and the propensity of all the conformations to convert to T is inadvertently the highest. Additionally, conversion to trident form is averted completely as is evident from the absence of that node in these networks.

We similarly, evaluated the inter-conversion of conformations for each TG in the droplet throughout the simulation (Supplementary Figure 13). It can be observed that the network formed in this case is a lot denser compared to that of a solvated single TG. Each node was observed to make connections with every other node unlike in Figure 3. The inclination towards a particular inter-conversion depicted by edge thickness may be dependent on the spatial location of TG in the droplet. In other words, the biasness towards a particular inter-conversion may be guided by the duration of exposure to water such that TGs on surface and those buried deeper in the core, may behave differently.

Next, we attempted to elucidate the mechanism of this inter-conversion by computing the free energy values of each of the six conformations,in both the medias. The probability distribution of all the molecular arrangements that existed in our system were evaluated using root mean square deviation (RMSD) and radius of gyration (Rg) (See Methods for more details). The free energy distribution profiles so obtained have been depicted in Figure 4. For long TG in water free energy profiles of stacker, trident and hand were found to be distinct from that of chair, fork and T, such that the former three had lower free energy. Additionally, each of these six conformations had a wide range between which their free energy was distributed. (Figure 4A). While this profile was unable to explain why stacker is the preferred intermediate for most conversions, despite trident and hand also having a lower free energy value, it did highlight a peculiar trend. It can be observed that the inter-conversion is guided by the need to attain a lower free energy. For instance, Chair form converts to T & stacker, both of which have lower free energy. A similar trend is observed for fork which also frequently converts to T & stacker. For medium and short TG energy distribution profiles in water showed that T, chair and fork have the lowest energy compared to the other conformations. Therefore, the highly frequent conversions that we observed in this case, between stacker → T, hand → T appear to be guided by a need to gain higher stability, by converting to a conformation with a lower free energy value (Figure 4B & C). In cyclohexane, a similar trend was observed where stacker and hand converted to T (Figure 4D-F). All the free energy profiles (except for that of long TG in water) also highlight that fork has maximum stability. Despite this, it is not the most favored conformation as is evident from its frequency (Figure 2) as well as pattern of inter-conversion (Figure 3). This may also be a result of the choice of reaction coordinate used to define the molecular arrangements in our system. As shown in Supplementary Figure 14-16, while RMSD and Rg values do vary for each of the six conformations, they do not yield distinct clusters.

**Figure 4:**
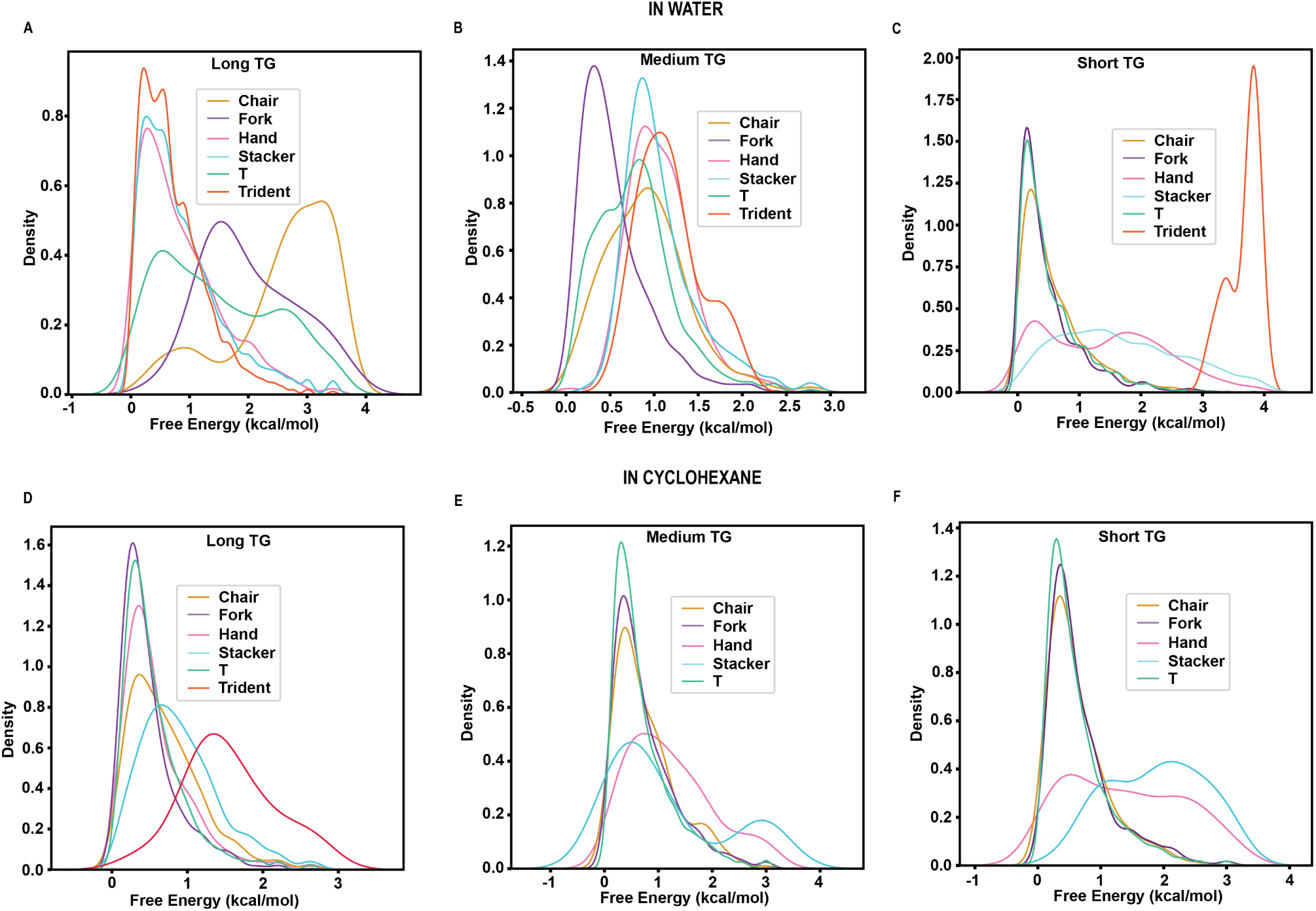
Free energy distribution of TG in water and cyclohexane. The free energy calculated based on root mean square deviation (RMSD) and radius of gyration (Rg) of each of the six conformations throughout the simulation has been plotted. Panel A,B & C show energy distribution of long, medium & short TG in water, respectively. Panel D, E & F show energy distribution profile of long, medium & short TG in cyclohexane, respectively.

Fatty acids encompass a heterogenous class that vary in their length of acyl chain carbons and number of unsaturation. Particularly, fatty acids with *>* 16 carbons, may be saturated or unsaturated. Depending on the level of unsaturation, they may confer unique physical properties such as elasticity and interdigitation, which in turn influence the structural organization and solvation properties ^26;13^. Therefore, we took a case of long TG (C18) and studied the effect of unsaturation on its solvation properties and structural organization. We observed that the presence of C = C, does not alter the hydrophobic property of acyl chain drastically. This is evident from the exactly identical *g(r)* profile of water around COM of acyl tail (Supplementary Figure 17 A-B). Infact, there were only minor differences observed in the ΔG_solv_ of the two long TGs (Supplementary Figure 17C). Next, we evaluated if the presence of unsaturation offers a predisposition towards any particular conformation. We observed no difference in case of a TG droplet, where both the droplet made of only saturated long TG and that made of only unsaturated TG, existed in the stacker form (Supplementary Figure 18A). This may also indicate that this form has an efficient packing tendency. However, when a single TG was immersed in water, the presence of unsaturation shifted the preferred conformation from trident to stacker (Supplementary Figure 18B). This is likely because the presence of a double bond increases the rigidity of the acyl tail, limiting its degree of freedom in water.

## Conclusion

In summary, we have elucidated the solvated structure of an isolated TG and confirmed its preference for hydrophobic environment based on its ΔG_solv_ in cyclohexane and water. When immersed in water, TG forms H-bonds using its carbonyl oxygen, bringing a few water molecules closer while the acyl tails, together exert a dominating hydrophobic effect that keep the bulk water at a distance. This hydrophobic effect is dependent on the length of acyl chains, such that a long chain hydrocarbon exerts a stronger force. Highly positive values of ΔG_solv_ in water are also indicative of the inability of this non-polar solute to overcome the H-bonding of water. We have also shown that a single TG molecule flips between all the six conformations during solvation, but its preference for a particular conformation is guided by the length of its acyl tails and the polarity of solvent. Long TG prefers to exist as trident in water while medium and short TG exist as T in water. In cyclohexane, irrespective of the length of acyl chain, all the three TGs exist as T. The presence of unsaturation was also observed to affect this preference, causing a shift from trident to stacker in case of long unsaturated TG. These preferences are in part, a result of the comparatively lower free energy values of each of these conformations, in the respective solvent.

In a lipid droplet constituted by only long TG, stacker form is the predominant form, highlighting its efficiency in packing. The inter-conversion between the six forms is guided by the need to exist in a conformation which is more stable in the medium. In case of water, a compact conformation like stacker or trident is stable because the acyl tails are closer to each other which is followed by hand, chair and fork. T represents a conformation where the acyl tails are most extended and therefore, this conformation is only stable in a non-polar medium, where these chains are able to form strong stabilizing van der Walls interactions with the solvent. This also explains why this conformation is the prevalent conformation in the bulk lipid droplet ^13^. To conclude, this study has provided an atomic description of the solvation process of an isolated TG while simultaneously highlighting similarities and differences to its structural organization in the bulk. We believe this work will pave way to future computational studies that may shed light on the molecular description of the process of lipid transfer and exchange.

## Acknowledgment

The authors thank Prof. Ilpo Vattuleinen (University of Helsinki) and Patrick Fuchs (Paris Diderot University) for providing the GROMOS parameters of triolein. We also thank Amélie Bacle (Institute Jacques Monod - CNRS University Paris City), Stefanno Vanni (University of Fribourg) for their help in classifying the triglycerides into various conformations. We would like to also thank Sriram K and Arul Murugan (Indraprastha Institute of Information Technology, Delhi) for their insightful inputs and discussions.

