## Supplementary file for "Solvation Dynamics of a Single Triglyceride as a Function of its Chain Length"

### Experimental Methods

#### Atomistic Simulations

MD simulations were performed in triplicates with the GROMACS 2021.1 suite.[1–3] The system consisting of a single triglyceride (either long, medium or short) was represented by GROMOS96 53A6 force field, extended to include Berger lipid parameters (JCC 2004 vol 25 page 1656).[4,5] The water was modeled using SPC representation.[6] Cyclohexane was represented by GROMOS96 53A6 force field generated using PRODRG.[7] Each of the three starting conformations were placed in a cubic box large enough to contain the system with at least 1.0 nm of solvent on all sides. Periodic boundary conditions were used , and the long-range electrostatic interactions were treated with the particle mesh Ewald method[8] using a grid spacing of 0.16 nm (for water simulations) and 0.12nm (for cyclohexane simulations) combined with a fourth-order B-spline interpolation to compute the potential and forces in-between grid points. The real space cutoff distance was set to 1.4 nm and the Van der Waals cutoff to 1.4 nm. The bond lengths were fixed,[9] and a time step of 2 fs for numerical integration of the equations of motion was used. Coordinates were saved every 2 ps. The pressure coupling was done by employing a Berendsen barostat[10] using a 1 bar reference pressure and a time constant of 2.0 ps with compressibility of 4.5e-5 bar using isotropic scaling scheme. In case of cyclohexane, a time constant of 0.5 ps with compressibility 1.21e-8 bar[11] using isotropic scaling scheme was used. In case of lipid droplets, a single TG (either long, medium or short) was randomly placed in a box until an average density of 1030 kg/m3 was reached[5]. All the starting structures were subjected to a minimization protocol for 50000 steps using the steepest descent method followed by equilibration with restraints on lipid for 1 ns. Systems with cyclohexane were equilibrated for 10 ns to obtain an average density of ≈760 kg/m3.[12] Systems with lipid droplets were also equilibrated for 10 ns to obtain a stable system. Three independent trajectories, each of 50 ns at 310K were carried out in case of single TG in water or cyclohexane. While the lipid droplets were simulated for 100ns at 310K.

#### Classification of TG conformations

Triglycerides were classified according to the classification proposed by Backle et al.[13] in their work. The criterion and methodology for classification was the same as their work. Briefly, three unit vectors between each glycerol carbon upto the eighth last carbon in each fatty acyl chain were defined. In case of long TG [C1 to C12 Supplementary Figure 1 & 6]. Alternatively, for medium TG, the vectors were defined till the fifth last carbon of the fatty acyl chain [C1 to C7]. For short TG, vectors between the three glycerol carbons as well as the last carbon of the fatty acyl chain were considered [C1 to C5]. In case of long and medium TG, the entire length of fatty acyl chain was not considered to define the vector to account for the huge fluctuations in the terminal part of these chains (Supplementary Figure 6).Three scalar products that defined the relative orientation of each chain pair were computed. These products for each conformation of long, medium and short TG are shown in Supplementary Figure 7-9. For each of the seven conformations, namely, 'Trident', 'Fork', 'Chair', 'T', 'Right-Hand', 'Stacker', an ideal scalar product was defined. Next, for each scalar product in the sample space, the euclidean distance (d) from the ideal scalar product was computed. If the euclidean distance (d) $< 0.7$, the TG was assigned a conformation or else classified as 'Other'.

Each frame in the simulation was similarly assigned a conformation and the sum of each conformation at the end of the simulation was used as a measure of prevalence of a particular conformation in aqueous solution. In case of lipid droplet, each lipid in each frame was assigned a particular conformation.

#### Free energy distribution for each conformation of triglyceride

Free energy for each conformation was computed based on the following formulae :

$$\Delta G(R) = -k_B T \; [ln(P(R)) - ln(P_{max})] \tag{1}$$

where $k_B$ is the Boltzmann constant, P is the probability distribution of the molecular system along some coordinate R, and $P_{max}$ denotes its maximum, which is subtracted to ensure that $\delta G = 0$ for the lowest free energy minimum. R is the order parameter of the system and was chosen to be the range of root mean square deviation (RMSD) as well as radius of gyration (Rg) values for each of the triglyceride conformation.

#### Inter-conversion of triglyceride conformations

Each of the six conformations of a triglycerides represent the six states that it can exist in and their inter-conversion has been depicted as a network. Simulation trajectories were analyzed to compute the succeeding conformation for a given state. For instance, the absolute values obtained represent the number of times a stacker was succeeded by a trident. For each such conformation, the absolute values were normalized using min-max normalization as follows:

$$x' = \frac{x - min(x)}{max(x) - min(x)} \tag{2}$$

These normalized values were amplified by multiplying by a factor of 10, and the values obtained were taken as is to define the edge weights in the network. Briefly, the network is composed of six nodes, each of which define the six states or conformation a triglyceride can exist in. The size of the node is determined by the number of connections it makes with the other nodes. Connections between two nodes are defined as edges and are directional in nature. The direction of the arrow points towards the conformation after conversion.

#### Number of water molecules surrounding a conformation

The number of water molecules around each conformation were calculated using gmx trjorder by defining a shell radius of 0.5nm.

#### Radial distribution function

Radial distribution function (RDF) defined as follows:

$$g(r) = \frac{\rho(r)}{\rho_0} \tag{3}$$

where $\rho$(r) is the number density at a particular distance r away from some specified atom and $\rho_0$ is the bulk density of the solvent. RDF was calculated using gmx rdf by defining the reference point as center of mass of triglyceride, center of mass of glycerol group, center of mass of acyl tails respectively.

#### Solvation free energy calculations

The solvation free energy of systems solvated in either water or cyclohexane was computed as the difference in free energy by transforming between two states (solvated and unsolvated) using a coupling parameter, ($\lambda$), approach in conjunction with thermodynamic integration (TI)[14,15] defined as:.

$$\Delta F_{AB} = \int_{\lambda_A}^{\lambda_B} \left\langle \frac{\partial H(\lambda)}{\partial \lambda} \right\rangle_{\lambda'} d\lambda' \tag{4}$$

In this approach the $\lambda$-dependence of the Hamiltonian $H$ is a function of the coupling parameter $\lambda$ and defines a pathway that connects the two states of the system denoted as 'State A' and 'State B' (solvated and unsolvated in this case). We evaluated the ensemble average at a number of discrete $\lambda$-points by performing separate simulations for each chosen $\lambda$-point. The integral was then determined numerically.

Generally speaking, the solvation free energy is defined as the sum of energies required to introduce a solute in the solvent; but in this case, the solvation free energy is calculated by decoupling the solute from the solvent according to the thermodynamic cycle shown above. $\Delta F_1$ is the work required to remove all the internal non-bonded interactions in the lipid in the vacuum. This is achieved by gradually mutating all atoms in a given state (State A) into 'dummy' atoms (State B). Therefore, a dummy atom is an atom for which non-bonded interaction, namely Lennard-Jones and electrostatic parameters, with all other atoms have been set to zero. The bonded interactions within the molecule as well as the masses of the individual atoms are kept unchanged. $\Delta F_2$ is the work required to transfer this dummy solute from vacuum to the solvated phase. As the molecule does not interact with the rest of the system, and the available volume is the same, this term is effectively zero. $\Delta F_3$ is the work required to remove the solute– solvent and the solute–intramolecular interactions. This is again achieved by gradually mutating all atoms in a given compound (state A) into "dummy" atoms (state B). Since the van der Waals interactions are gradually turned off between the solute and solvent by mutating it into a dummy atom, the solute is slowly decoupled from the solvent.[14,15] This thermodynamic cycle is summarized in the following illustration.

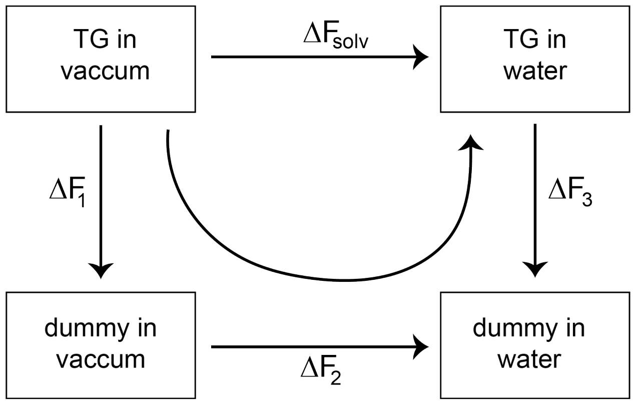


The TI method relies on $\lambda$ dependence of the Hamiltonian, $H$ to the solute-solvent interactions that gradually varies between full interactions ($\lambda = 0$) to no interactions ($\lambda = 1$). Therefore, the solute molecule was gradually made to disappear from the solution using the coupling parameter $\lambda$. For each $\lambda$ value, a series of independent molecular dynamics simulations were performed and the gradient of the Hamiltonian, $H$ as a function of coupling parameter $\lambda$ was averaged over a large number of equilibrated configurations obtained during the simulation which were numerically integrated to yield solvation free energy $\Delta G_{solv}$.

In the current study, the solute was a triglyceride molecule which is hydrophobic in nature described by GROMOS96 53A force field; therefore, only van der Waals interactions were considered for the computation of solvation free energy utilizing only the Lennard Jones potentials derived from GROMOS96 53A6 parameters. The electrostatic interactions were not taken into account, and therefore the charges were turned off during free -energy simulations.

The free energy simulation protocol utilized 20 $\lambda$ points spaced every 0.05 from 0 to 1. At each lambda point, the system was simulated for 1 ns at an isothermal-isobaric ensemble, providing values for $\Delta G_{solv}$. These free energy simulations were carried out for all three TGs in two solvents (water and cyclohexane) and prior to the simulations at different $\lambda$ points, each system was minimized using steepest descent algorithm. Once minimized, each system was subjected to an equilibration for 100 ps using a time step of 2 fs at the canonical and isothermal-isobaric ensembles. Solute-solvent non-bonded interactions were treated using the particle mesh Ewald method setting the cutoff range of 1.2 nm for Coulombics whereas the cutoff value for the evaluation of van der Waals interactions was also set to 1.2 nm. Temperature was controlled at 310K using a Langevin dynamics integrator with a coupling time constant of 1.0 ps. The pressure was kept fixed at 1 bar using the Parinello-rahman barostat[16] with a coupling time constant of 1.0 ps and compressibility value of 4.5 e-5 bar for water and 0.5 ps and 1.21 e-8 bar for cyclohexane.[11]

All free energy simulations were performed using GROMACS 2021.1, whereas computations of energy difference between the two states were carried out using the Bennett Acceptance Ratio (BAR) module of the GROMACS program.[17]

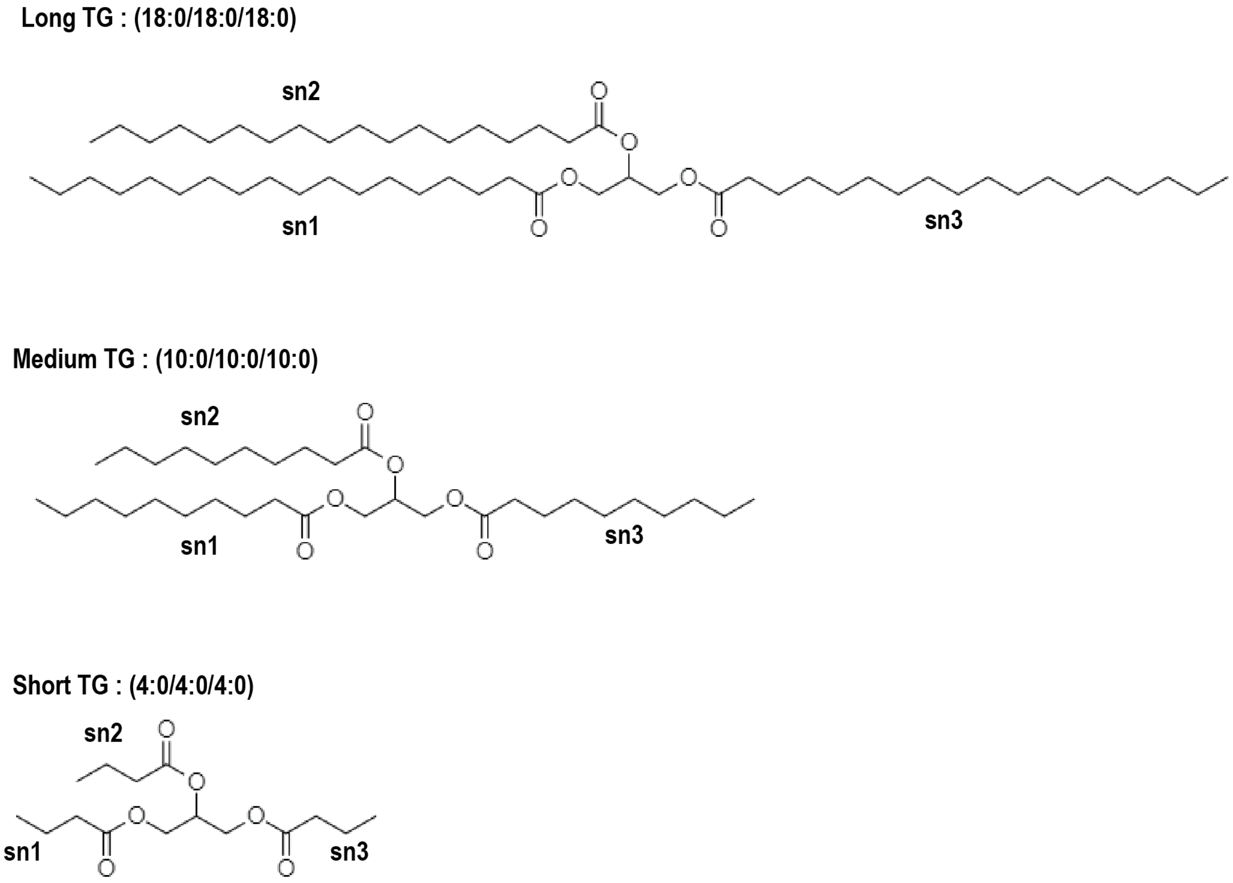


Figure 1: **The three TG structures varying in acyl chain length used in the study.**

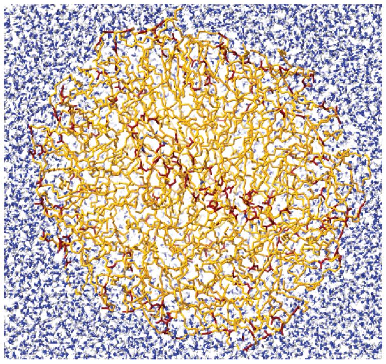

| Long droplet (18:0) | | @ 310 K |
|---|---|---|
| # of lipid molecules | # of water molecules | Total atoms (system size) |
| 108 | 71,656 | 221,772 |

| Medium droplet (10:0) | | @ 310 K |
|---|---|---|
| # of lipid molecules | # of water molecules | Total atoms (system size) |
| 92 | 24,354 | 76,650 |

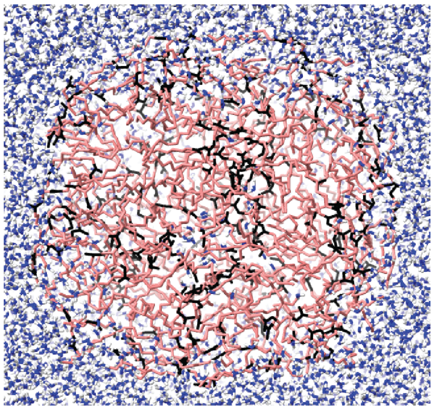

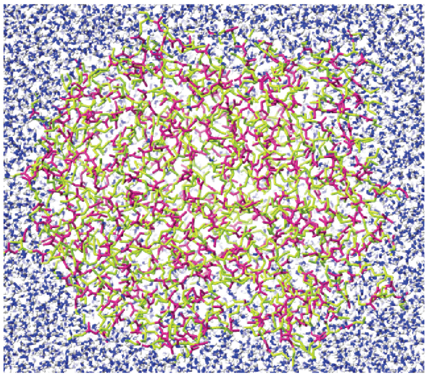

| Short droplet (4:0) | | @ 310 K |
|---|---|---|
| # of lipid molecules | # of water molecules | Total atoms (system size) |
| 282 | 38,361 | 121,005 |

| Long droplet 2 (18:1) | | @ 310 K |
|---|---|---|
| # of lipid molecules | # of water molecules | Total atoms (system size) |
| 108 | 71,656 | 221,772 |

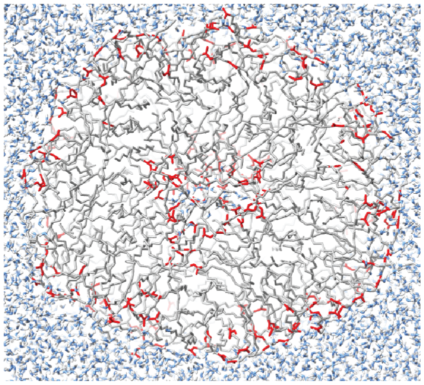

Figure 2: **100ns snapshots of the lipid droplet simulations, along with details of the systems.**

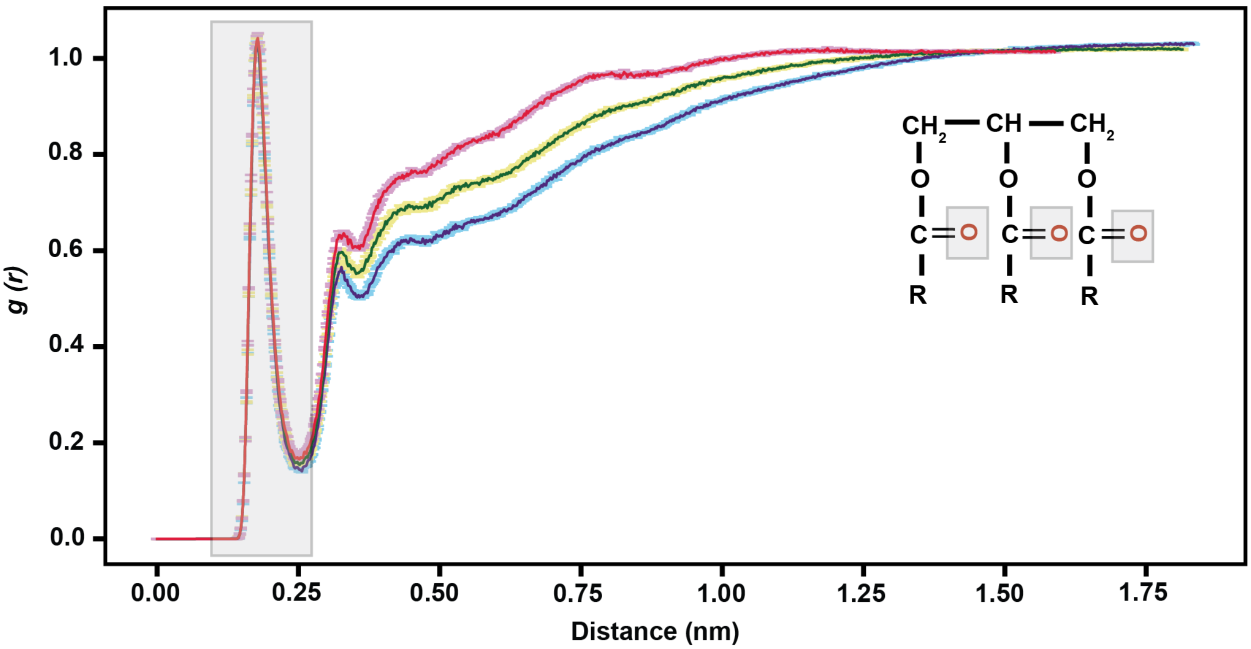


Figure 3: **Radial distribution profile for the hydrogen atoms of water around carbonyl oxygen of TG.** Solid line represents mean while error bars depict absolute deviation from mean across three simulations. The carbonyl oxygen atoms considered in the analysis have been highlighted in red in the TG structure and corresponds to the peak at ≈ 0.24 nm which is highlighted in gray.

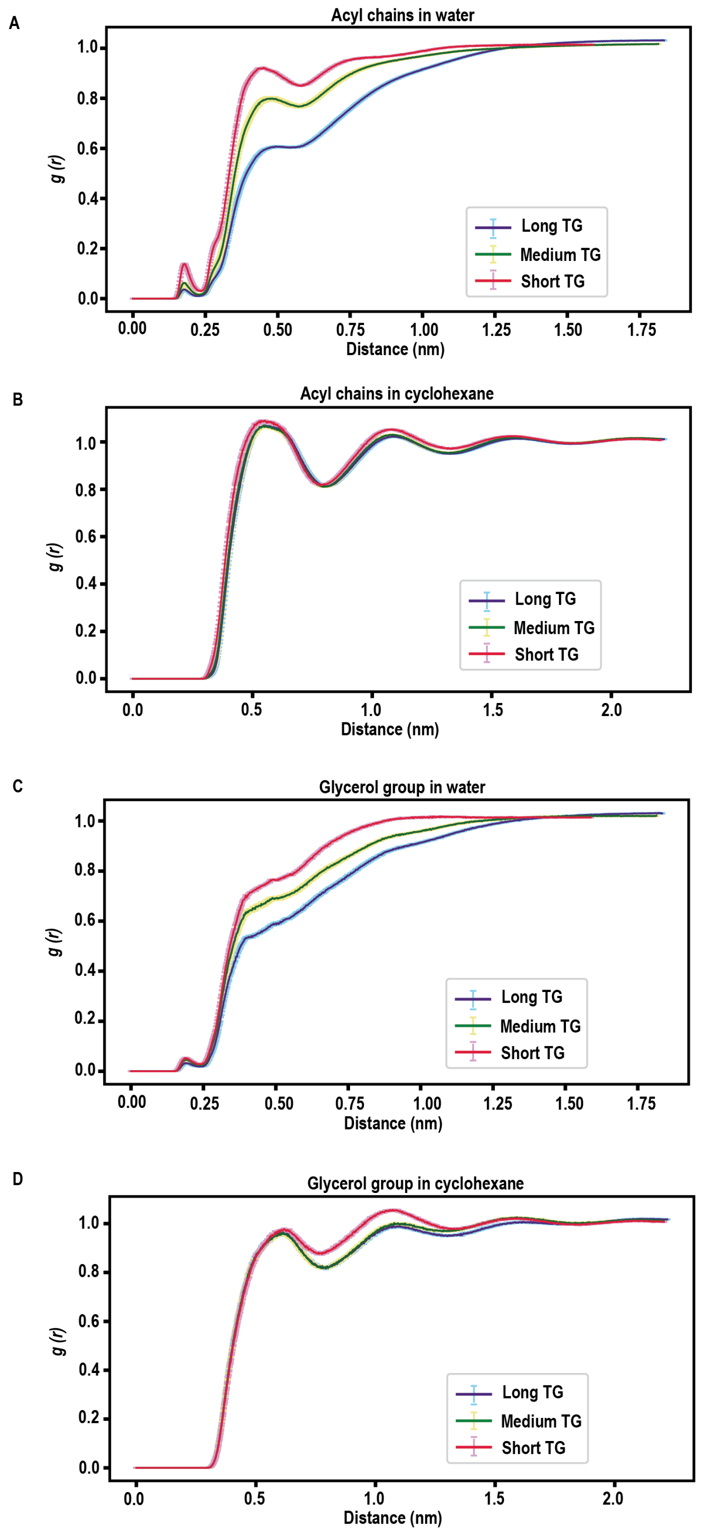


Figure 4: **Radial distribution profile of the solvent molecules around acyl chains as well as glycerol group of TG.** Solid line represents mean while error bars depict absolute deviation from mean across three simulations.

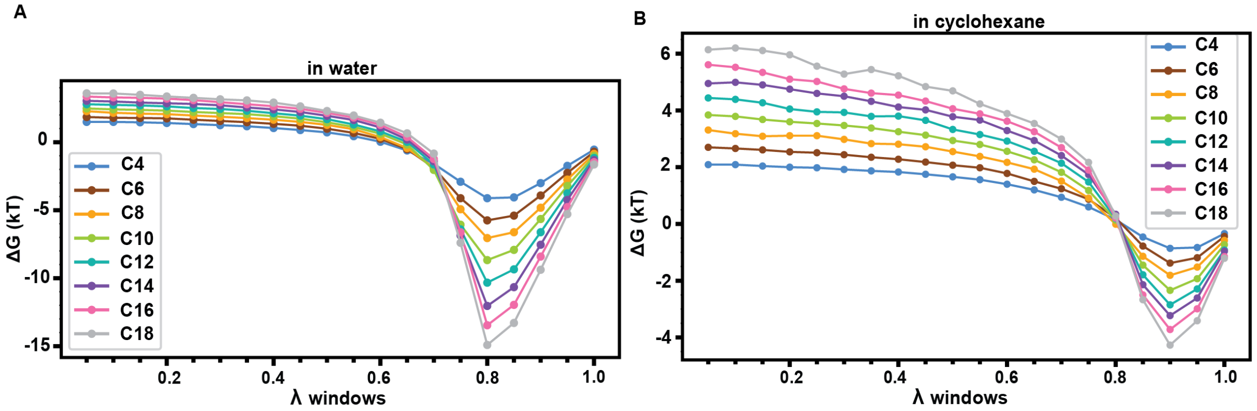


C

(in kJ $mol^{-1}$)

| | ΔG solv water | ΔG solv chx |
|---|---|---|
| C4 | 15.09 +/- 0.43 | -58.26 +/- 0.45 |
| C6 | 21.24 +/- 0.27 | -72.51 +/- 0.24 |
| C8 | 24.31 +/- 0.58 | -86.37 +/- 0.32 |
| C10 | 30.70 +/- 1.15 | -100.27 +/- 0.89 |
| C12 | 34.25 +/- 0.83 | -113.95 +/- 0.60 |
| C14 | 38.02 +/- 0.64 | -129.86 +/- 0.55 |
| C16 | 39.52 +/- 1.35 | -139.43 +/- 0.74 |
| C18 | 44.83 +/- 1.33 | -157.21 +/- 1.17 |

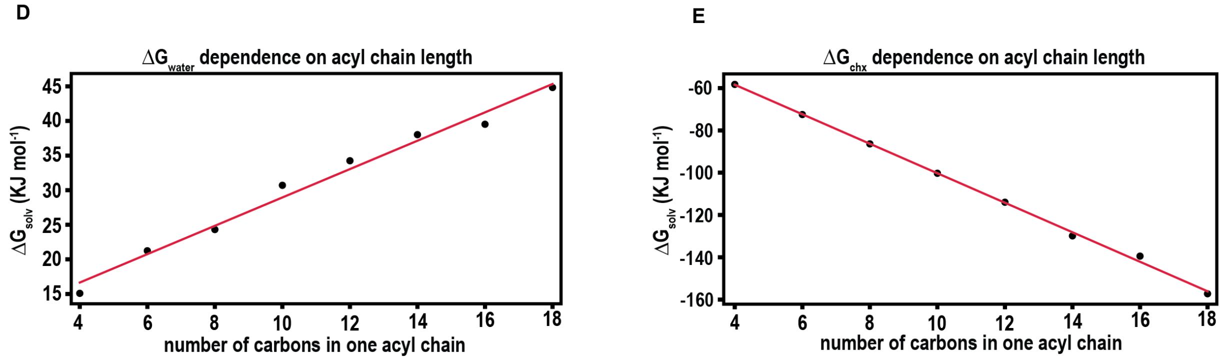


Figure 5: Free energy difference between neighboring values of $\lambda$ in **A.** water and **B.** cyclohexane for TGs with acyl chain lengths ranging between four to eighteen carbons. **C.** The free energy of solvation in water and cyclohexane for each of the eight TGs. The $\Delta G_{solv}$ dependence on acyl chain length has been shown in **D.** water ($R^2 = 98.39\%$) and **E.** cyclohexane ($R^2 = 99.85\%$).

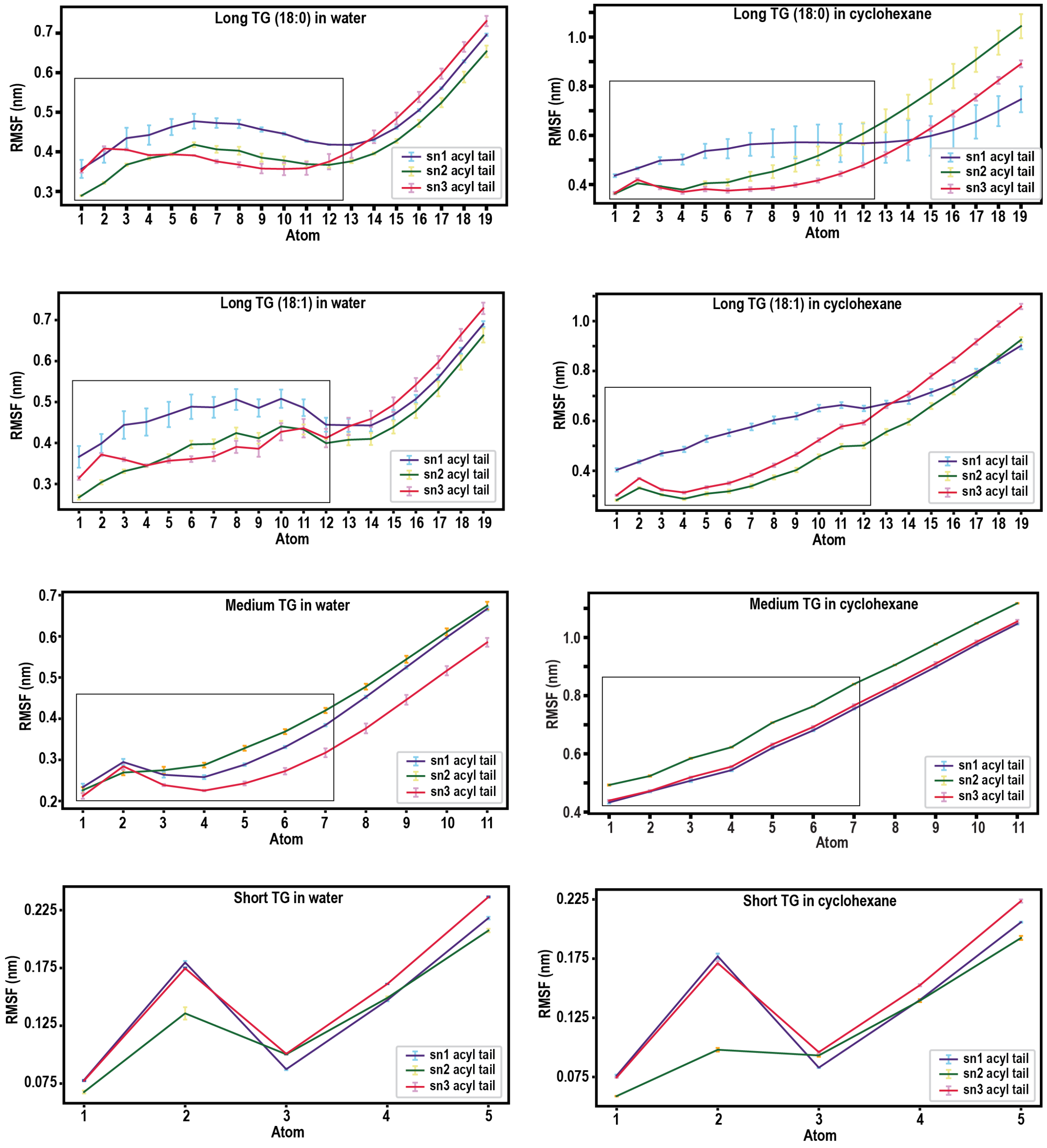


Figure 6: **Root mean square fluctuation (RMSF) of TG in water and cyclohexane.** The region in dotted line represents the region used for defining the vectors from the central carbon during assignment of conformations for both long and medium TG. In case of short TG, the terminal carbon of acyl chain was considered as mentioned in Materials and Methods.

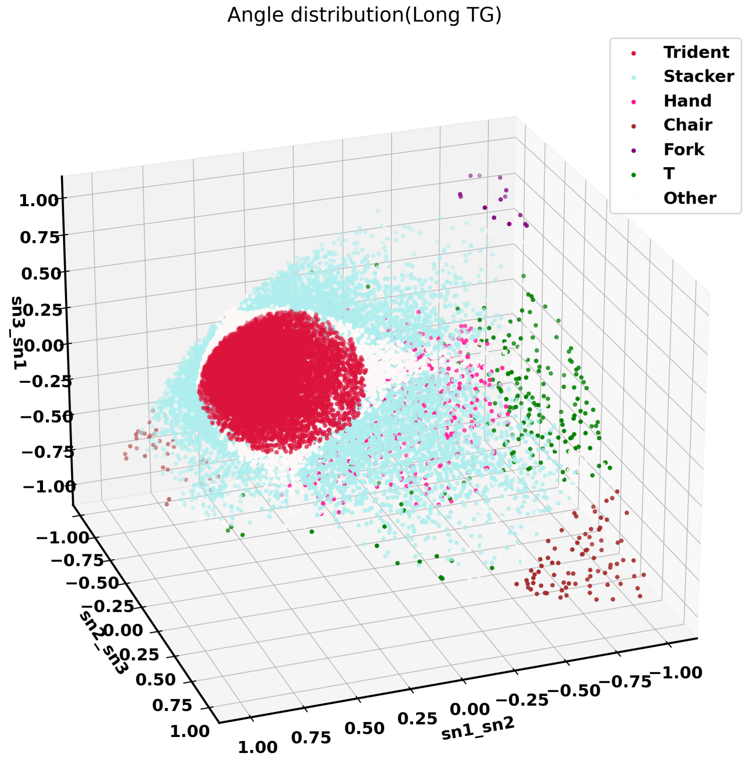


Figure 7: **Distribution of dot products between vectors of sn1, sn2 and sn3 chains used to classify the six long TG conformations.**

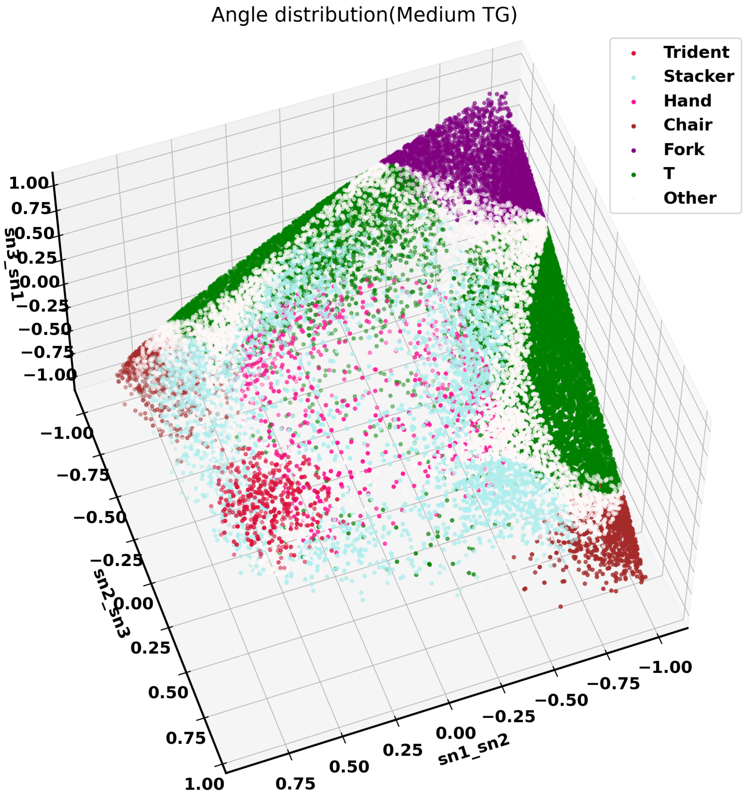


Figure 8: **Distribution of dot products between vectors of sn1, sn2 and sn3 chains used to classify the six medium TG conformations.**

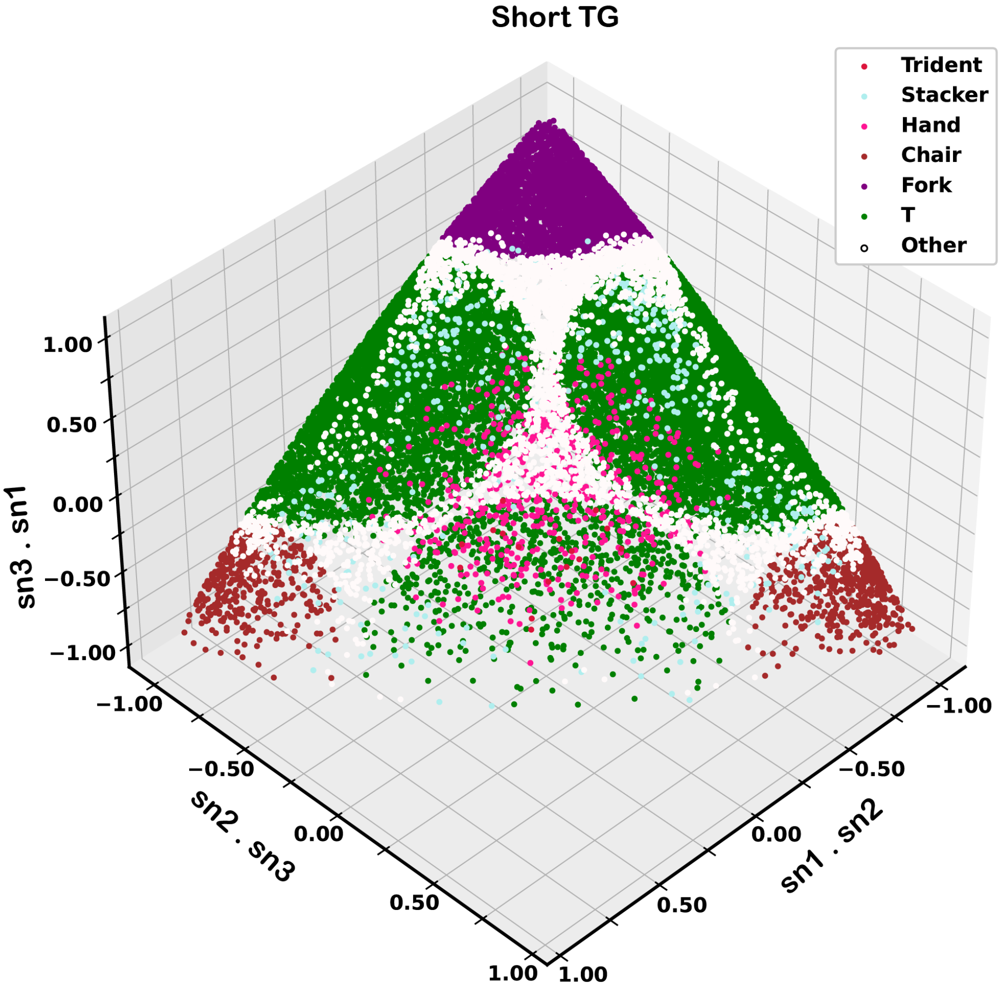


Figure 9: **Distribution of dot products between vectors of sn1, sn2 and sn3 chains used to classify the six medium TG conformations.**

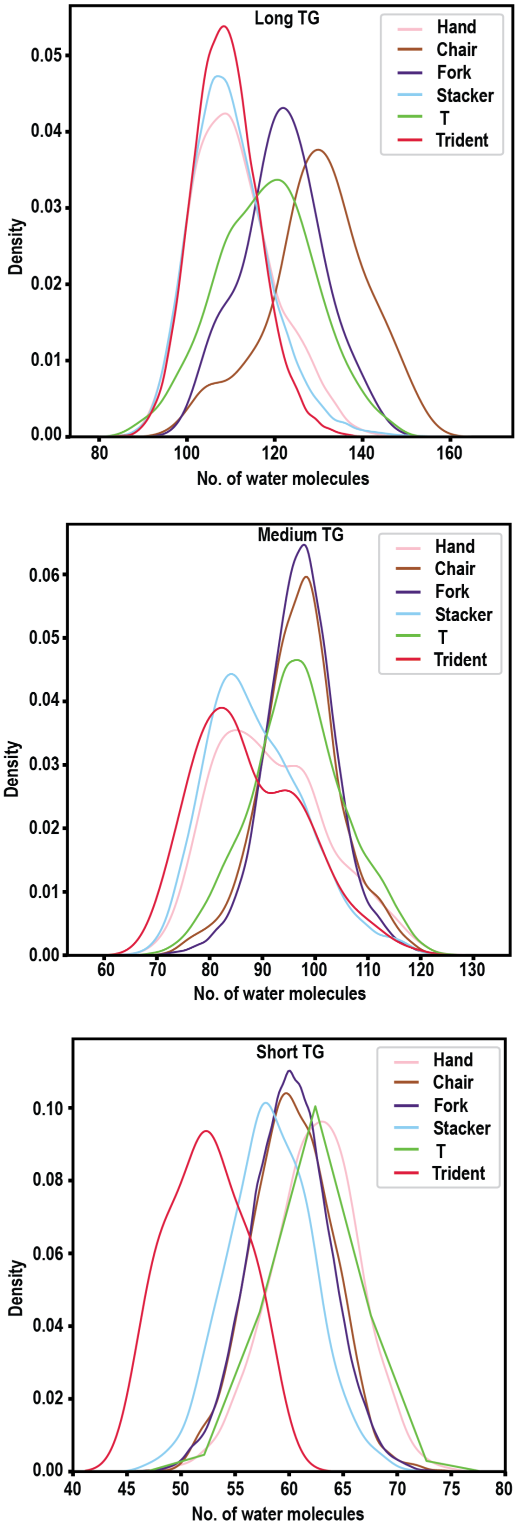


Figure 10: **Probability distribution curve of the number of water molecules within 0.5nm radius of the acyl tail for each of the six forms of long, medium and short TG**

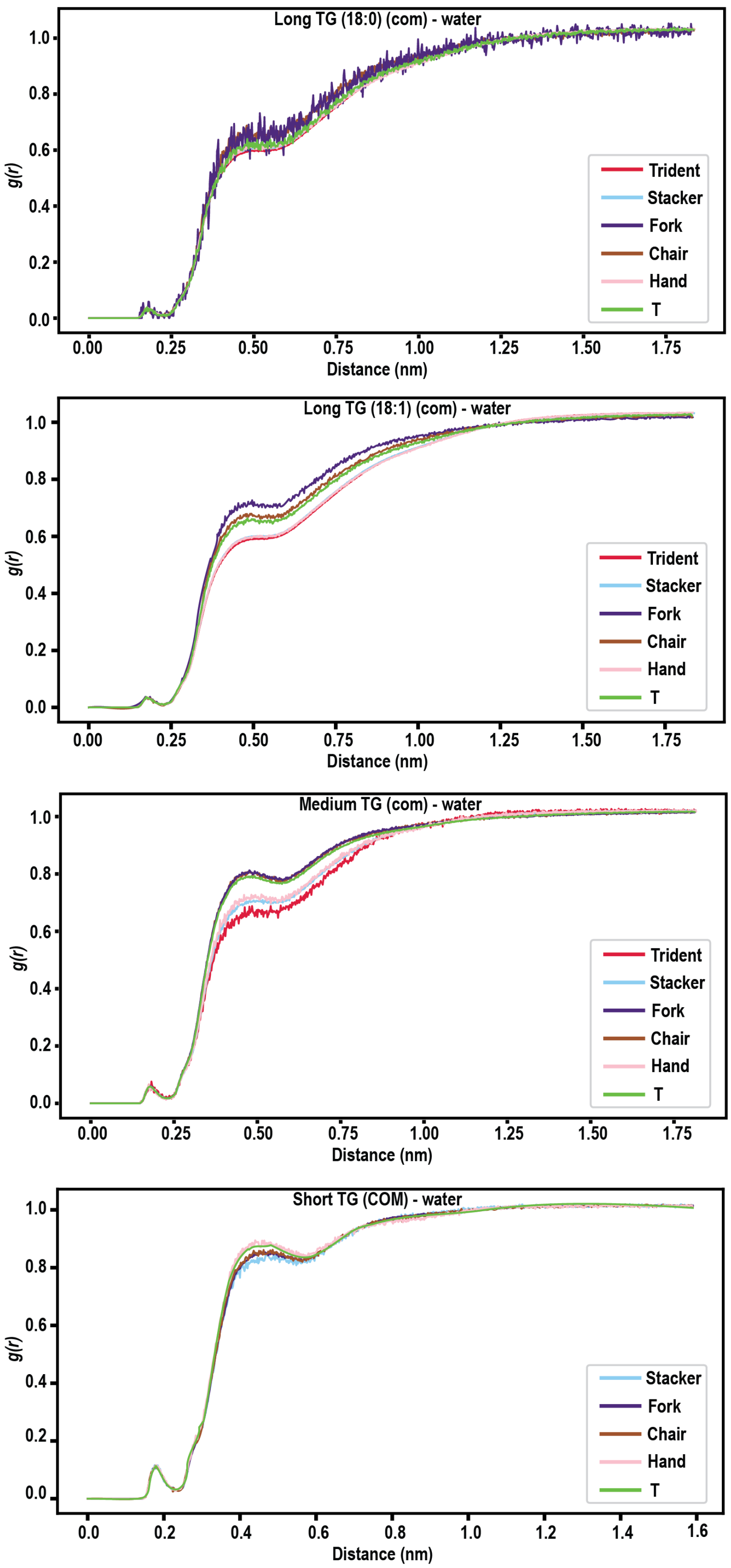


Figure 11: **Radial distribution profile of the COM of the six forms of each long, medium and short TG.**

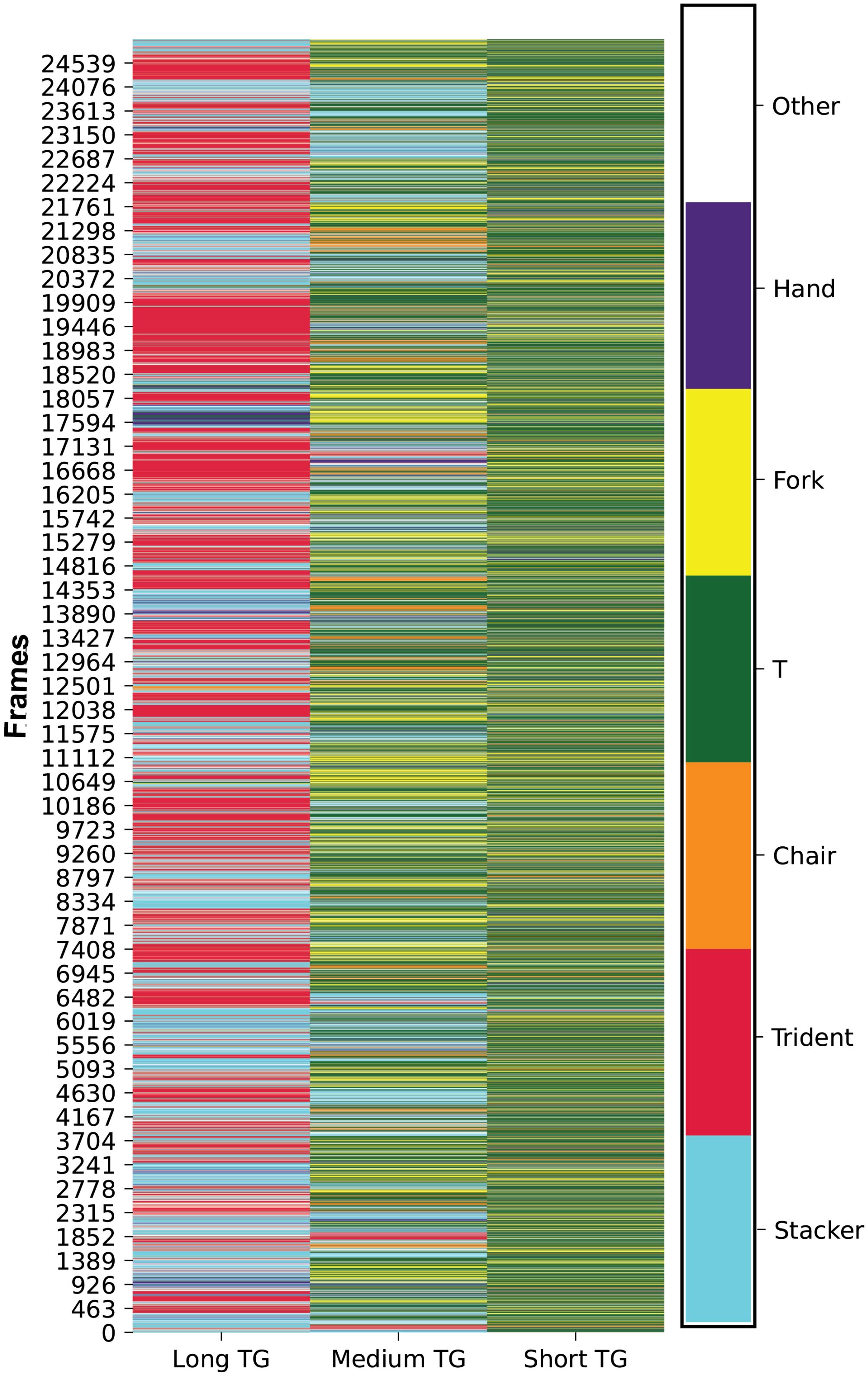


Figure 12: **Heatmap depicting the inter-conversion amongst the six forms throughout the simulation time for TG-water.**

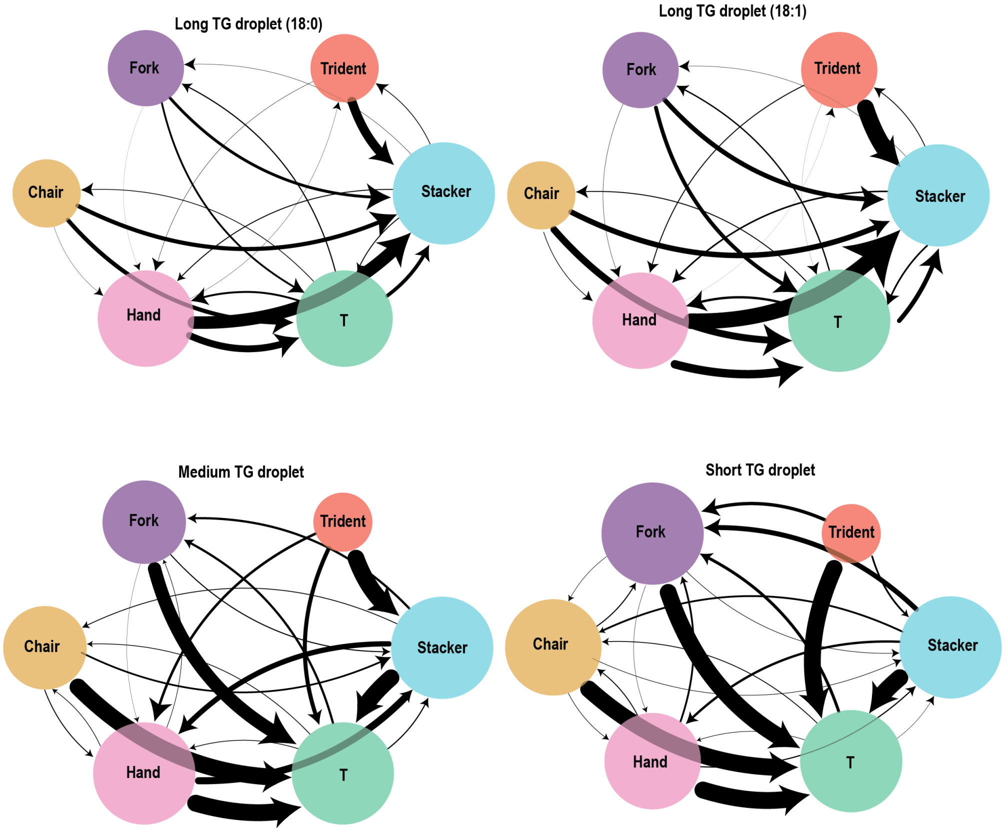


Figure 13: **Inter-conversion of triglyceride conformations of every TG constituting a lipid droplet.**A weighted network showing the inter-conversion between the six conformations. Each node in the network represents one of the six conformations as indicated in the figure and its size is proportional to the number of connections with other nodes. Nodes are connected to other nodes through edges which are directional such that the direction of arrow points towards the conformation after conversion. The thickness of the edges are proportional to the frequency of conversion observed between the two conformations that they connect.

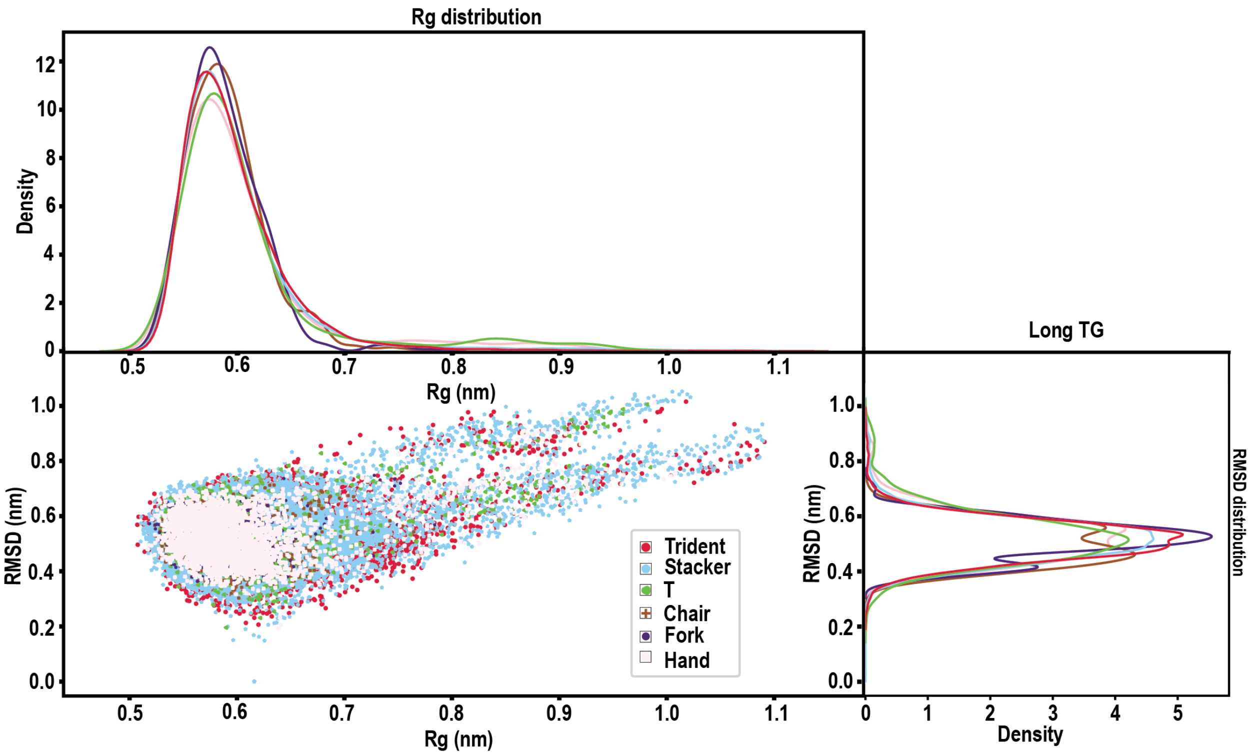


Figure 14: **Distribution of the RMSD as well as Rg values for each conformation of a long triglyceride.**

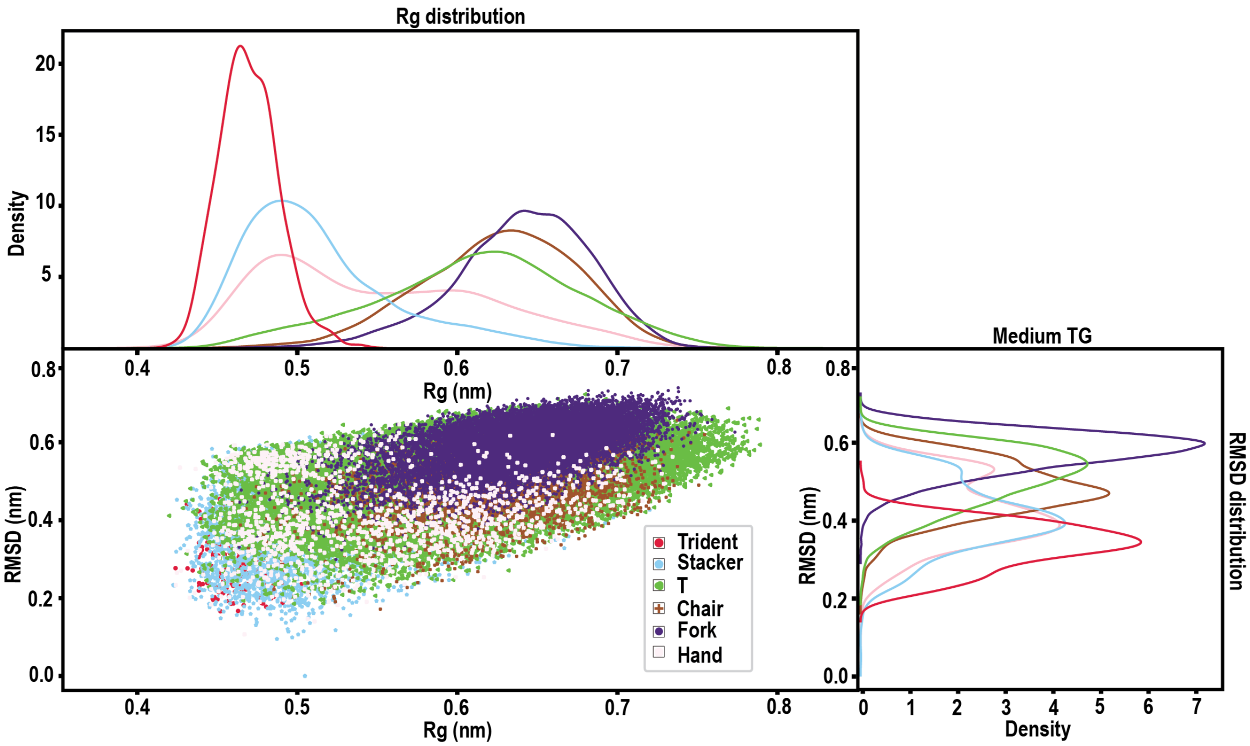


Figure 15: **Distribution of the RMSD as well as Rg values for each conformation of a medium triglyceride.**

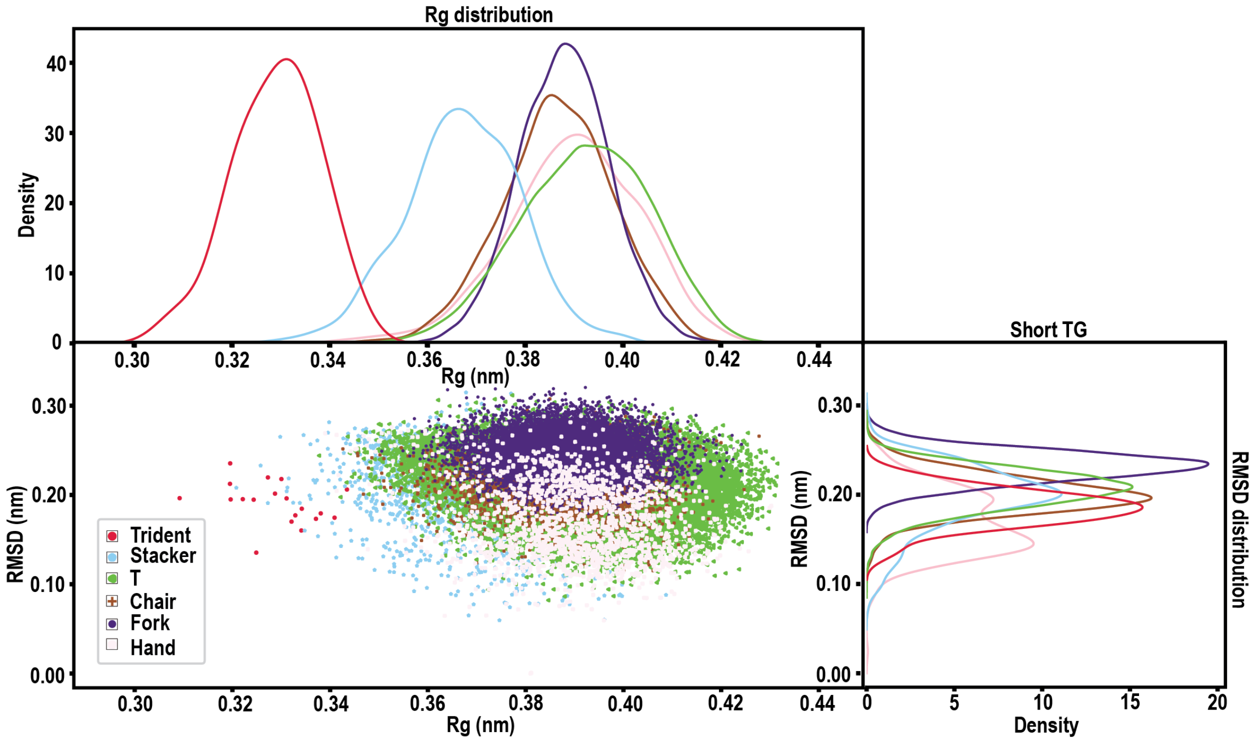


Figure 16: **Distribution of the RMSD as well as Rg values for each conformation of a short triglyceride.**

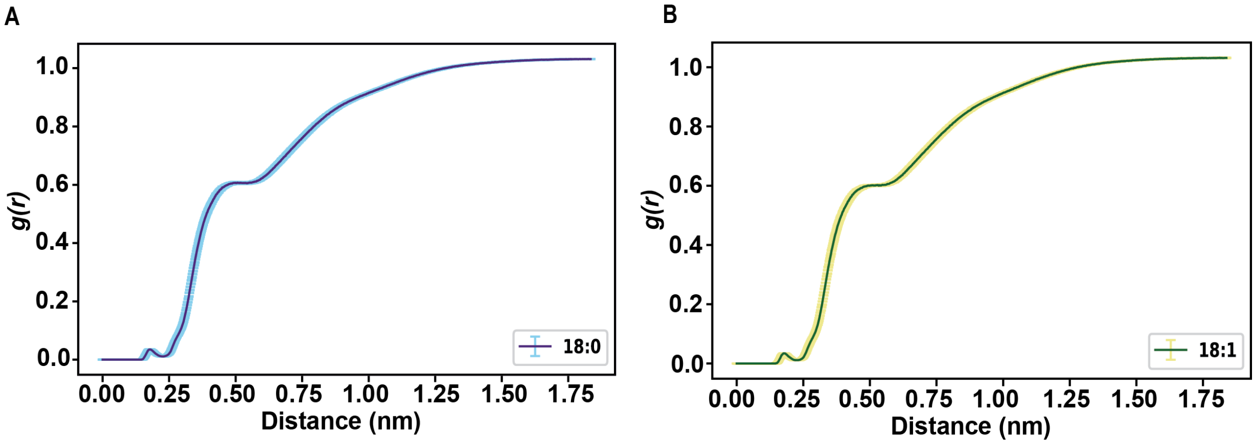


C

| (in kJ mol⁻¹) | 18:0 | 18:1 |
|---|---|---|
| ΔG solv water | 44.83 +/- 1.33 | 46.76 +/- 1.08 |
| ΔG solv chx | -157.21 +/- 1.17 | -143.93 +/- 1.41 |

Figure 17: **Effect of unsaturation of solvation properties of long TG.** Radial distribution profile of water molecules around the center of mass of acyl tails of **A.** long TG (18:0/18:0/18:0) **B.** long TG (18:1(9Z)/18:1(9Z)/18:1(9Z)) **C** Comparison of the free energy of solvation in water and cyclohexane of a single saturated and unsaturated TG.

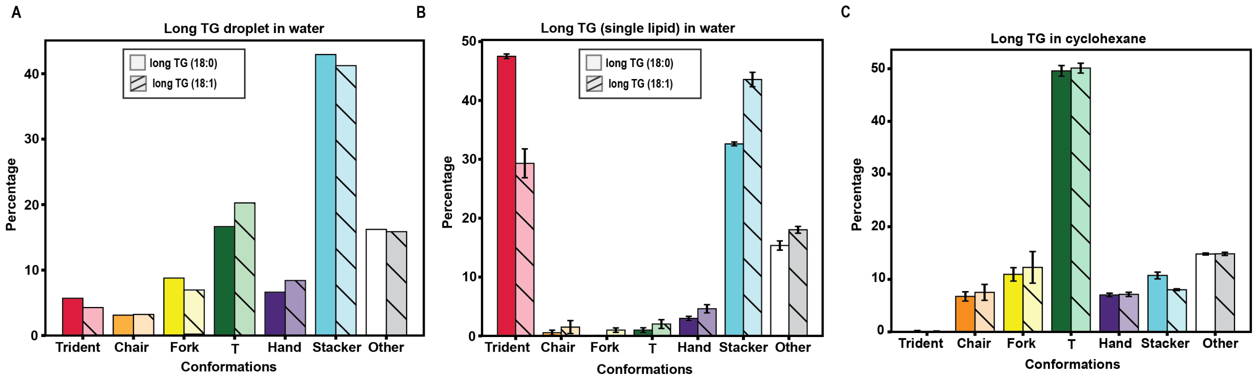


Figure 18: **Effect of unsaturation on propensity of TG towards a conformation.** The saturated long TG refers to (18:0/18:0/18:0) while unsaturated long TG refers to (18:1(9Z)/18:1(9Z)/18:1(9Z))
